# Interleukin-33 promotes type 1 cytokine expression via p38 MAPK in human natural killer cells

**DOI:** 10.1101/777847

**Authors:** David E. Ochayon, Ayad Ali, Pablo C. Alarcon, Durga Krishnamurthy, Leah C. Kottyan, Michael Borchers, Stephen N. Waggoner

**Affiliations:** Center for Autoimmune Genomics and Etiology, Cincinnati Children’s Hospital Medical Center, Cincinnati, OH, USA; Medical Scientist Training Program, University of Cincinnati College of Medicine, Cincinnati, OH, USA; Graduate Program in Immunology, University of Cincinnati College of Medicine, Cincinnati, OH, USA; Department of Pediatrics, University of Cincinnati College of Medicine, Cincinnati, OH, USA; Division of Allergy and Immunology, Cincinnati Children’s Hospital Medical Center, Cincinnati, OH, USA; Division of Pulmonary, Critical Care, and Sleep Medicine, University of Cincinnati College of Medicine, Cincinnati, OH, USA; Department of Internal Medicine, University of Cincinnati College of Medicine, Cincinnati, OH, USA

**Keywords:** Innate lymphoid cell, p38 MAPK, IL-12, NK cell, IL-33, IFN-γ, GM-CSF, TNF, synergy, asthma, COPD

## Abstract

This study tests the hypothesis that activation of mitogen-activated protein kinase (MAPK) by physiologically-relevant concentrations of interleukin-33 (IL-33) contributes to enhanced cytokine expression by IL-12 stimulated human natural killer (NK) cells. While IL-33 canonically triggers type 2 cytokine responses, this cytokine can also synergize with type 1 cytokines like IL-12 to provoke interferon-gamma (IFN-γ). We show that picogram concentrations of IL-12 and IL-33 are sufficient to promote robust secretion of IFN-γ by human NK cells that greatly exceeds responses to either cytokine alone. Nanogram doses of IL-33, potentially consistent with levels in tissue microenvironments, synergize with IL-12 to induce secretion of additional cytokines, including tumor necrosis factor (TNF) and granulocyte-macrophage colony-stimulating factor (GM-CSF). IL-33-induced activation of the p38 MAPK pathway in human NK cells is crucial for enhanced release of IFN-γ and TNF in response to IL-12. Mechanistically, IL-33-induced p38 MAPK signaling enhances stability of *IFNG* transcripts and triggers ADAM17-mediated cleavage of TNF from the cell surface. These data support our hypothesis and suggest that altered sensitivity of NK cells to IL-12 in the presence of IL-33 may have important consequences in diseases associated with mixed cytokine milieus, like asthma and chronic obstructive pulmonary disease.

**Graphical Abstract:** 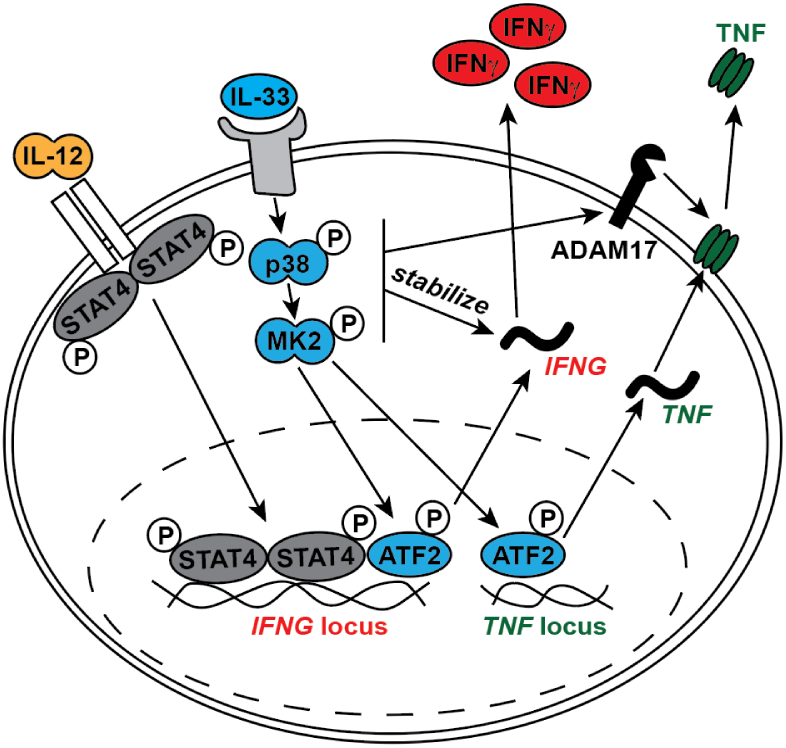

## Introduction

Natural killer (NK) cells play a key role in the clearance of virus-infected cells and the pathogenesis of many diseases^1-3^. The type 1 cytokine interleukin-12 (IL-12) provokes a canonical interferon-gamma (IFN-γ) response by NK cells^4^. Other type 1 cytokines, including IL-15, IL-18 and IL-1β enhance IL-12-induced IFN-γ release by NK cells^5-10^. This type 1 cytokine synergy can also promote enhanced release of tumor necrosis factor-alpha (TNF) and granulocyte macrophage colony stimulating factor (GM-CSF). Thus, type 1 cytokine rich environments associated with immune insults such as infection or injury can stimulate cytokine responses from NK cells that further promote type 1 responses. For example, in the experimental model of *Leishmania infantum*, the combination of IL-12 and IL-18 is critical for NK-cell IFN-γ expression^11^.

A central dogma of cytokine biology holds that type 1 cytokines (e.g. IL-12) suppress type 2 cytokine responses, while type 2 cytokines (e.g. IL-4) correspondingly suppress type 1 responses^12, 13^. Thus, type 2 cytokines should putatively suppress NK-cell production of IFN-γ. Yet the prototypical typical type 2 cytokine IL-4, alone or in combination with IL-12, triggered high levels of IFN-γ expression by mouse NK cells^14, 15^. Another type 2 cytokine, IL-33, can enhance IL-12-induced production of IFN-γ by both NK and NKT cells^16-18^. Thus, NK cells in type 2 cytokine rich environments may exhibit hypersensitive IFN-γ responses following IL-12-inducing infections or insults.

In the present study, we confirm that primary human NK cells treated with a combination of IL-33 and IL-12 *ex vivo* produce high levels of IFN-γ^16^, and we extend this observation to near physiological picogram concentrations of these cytokines. Mechanistically, we implicate the p38 mitogen-activated protein kinase (MAPK) pathway in IL-33-mediated enhancement of IL-12-induced responses of human NK cells and uncover a role for ADAM17 in enhanced release of TNF under these stimulatory conditions. These data provide mechanistic insights into how IL-33-rich inflammatory milieus may directly enhance proinflammatory activities of NK cells.

## Results

### IL-33 enhances IL-12-induced cytokine expression in primary human NK cells

IL-33 can bolster IFN-γ protein expression by IL-12 stimulated human NK cells^16^, but whether this enhancement occurs at level of transcription is unknown. To test this, we isolated NK cells from the blood of healthy de-identified adults prior to incubation of these cells in media containing IL-12, IL-33 or a combination of these cytokines for six hours. A concentration of 1 ng/mL of IL-12 induced a 10-fold increase of *IFNG* expression compared to unstimulated cells, while 0.5 ng/mL of IL-12 was insufficient to induce this response (**Fig. 1A**). The addition of IL-33 to NK cells cultured with either dose of IL-12 resulted in a >100-fold increase (Interaction: p<0.0001, two-way ANOVA) in *IFNG* mRNA expression levels (**Fig. 1A**). In a similar fashion, *TNF* transcript expression was increased ∼2.5-fold (Interaction: p=0.0061, two-way ANOVA) by the combination of IL-12 and IL-33 in comparison to IL-12 alone (**Fig. 1B**). In other types of innate lymphocytes, IL-33 can stimulate IL-5 and IL-13 expression^**19**^. In contrast, IL-33 alone or in combination with IL-12 had no measurable effect on expression of *IL5* or *IL13* expression by human NK cells (**Fig. 1C**).

**Figure 1.**
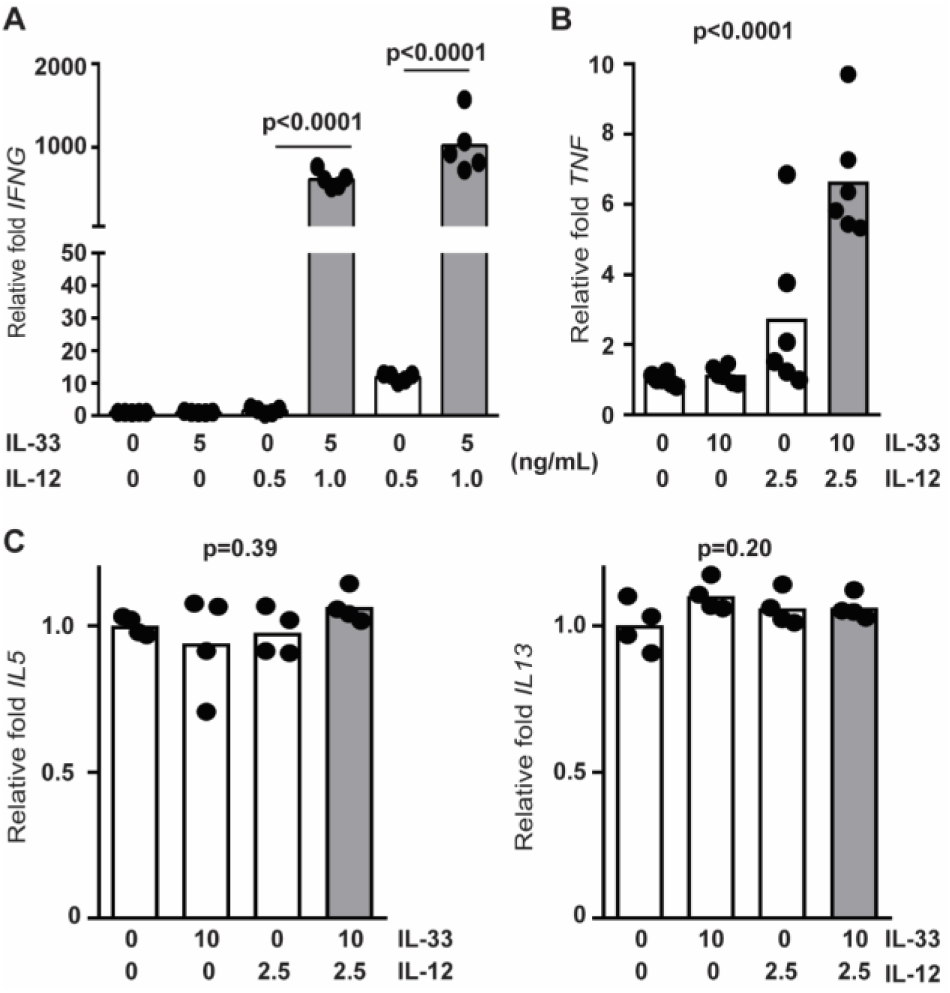
Increased *IFNG* and *TNF* expression in IL-12/IL-33 stimulated NK cells. Enriched primary human NK cells (4 to 6 different donors) were stimulated with combinations of IL-12 and IL-33 (doses listed in ng/mL) for 6 hours prior to qRT-PCR determination of (**A**) *IFNG*, (**B**) *TNF*, (**C**) *IL5*, and *IL13*. Signifance differnces in group means determined by one-way ANOVA (p values shown), while synergistic cytokine interactions assessed by two-way ANOVA are reported in the *Results* text. Each dot represents the mean of three biological replicates from an independent subject.

Next, we sought to examine whether near-physiological concentration of IL-33 could provoke enhanced IFN-γ expression in IL-12 stimulated NK cells. Isolated NK cells secreted IFN-γ in response to doses of IL-12 as low as 250 pg/mL (**Fig. 2A**). In contrast, production of IFN-γ by these cells was barely detectable after stimulation with IL-33 alone, even at doses as high as ≥1 ng/mL (**Fig. 2A**). However, 100 pg/mL or more of IL-33 enhanced (1.7- to 2.9-fold) IL-12-elicited IFN-γ protein expression (**Fig. 2A**), with synergistic interactions between IL-12 and IL-33 contributing significantly (p<0.0001, two-way ANOVA) to the overall variation in IFN-γ expression. High concentrations of IL-33 (10 ng/mL) additionally provoked expression of TNF and GM-CSF when administered in combination with IL-12 (**Fig. 2B**). The bulk of variation in the expression of TNF (p=0.011, two-way ANOVA) and GM-CSF (p=0.0008, two-way ANOVA) were attributable to synergistic interactions between IL-33 and IL-12 (**Fig. 2B**).

**Figure 2.**
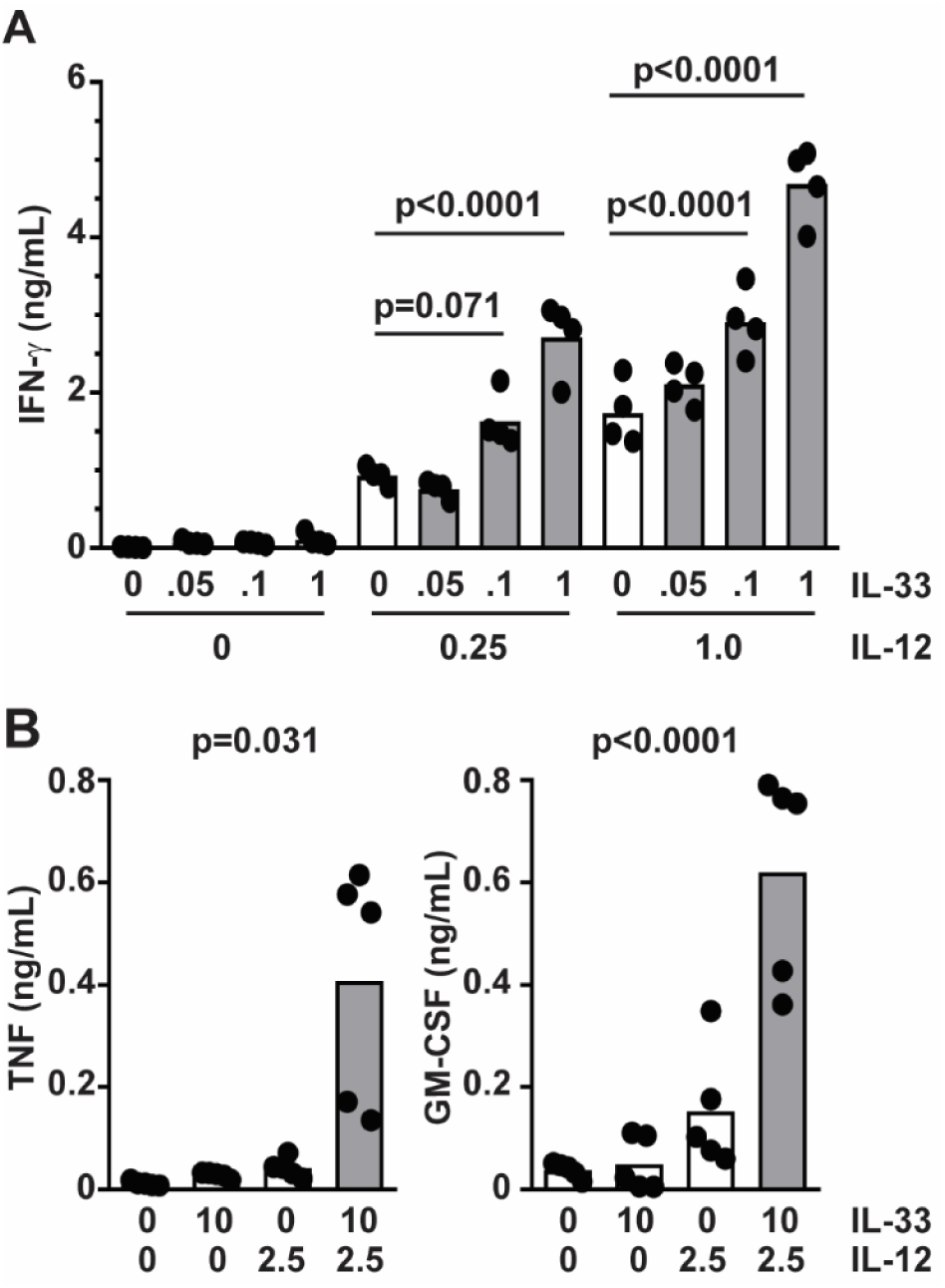
IL-33 enhances IL-12-induced cytokine expression in primary human NK cells. Enriched primary human NK cells were stimulated with combinations of IL-12 and IL-33 (doses listed in ng/mL) for 16 hours prior to ELISA determination of (**A**) IFN-γ, (**B**) TNF or GM-CSF expression in cell-free supernatant. Results from one representative iteration of (**A**) ten or (**B**) five similar experiments using cells from different subjects is shown (each dot is biological replicate and columns represent mean). Results of ordinary one-way ANOVA (**B**) assessment of group means with multiple comparisons (**A**) is shown. Two-way ANOVA was used to assess synergistic cytokine interactions (reported in the *Results* text).

Antibody blockade of the IL-33 receptor, ST2, abrogated the enhancing effect of IL-33 on IL-12-induced IFN-γ expression, consistent with previous studies^16^, but had no measurable effect on IFN-γ induced by IL-12 alone (**Fig. 3A**). Since IL-33 is part of the IL-1 superfamily that includes IL-18^20^, and IL-18 is a potent costimulator of IFN-γ expression by NK cells^9^, we sought to examine if the IL-18 receptor (IL-18R) plays any role in the synergy observed with IL-12 and IL-33. In contrast to ST2 blockade, IL-18R blockade had no effect on the ability of IL-33 to enhance IFN-γ release (**Fig. 3B**), but did prevent synergy of IL-18 with IL-12 (**Fig. 3C**).

**Figure 3.**
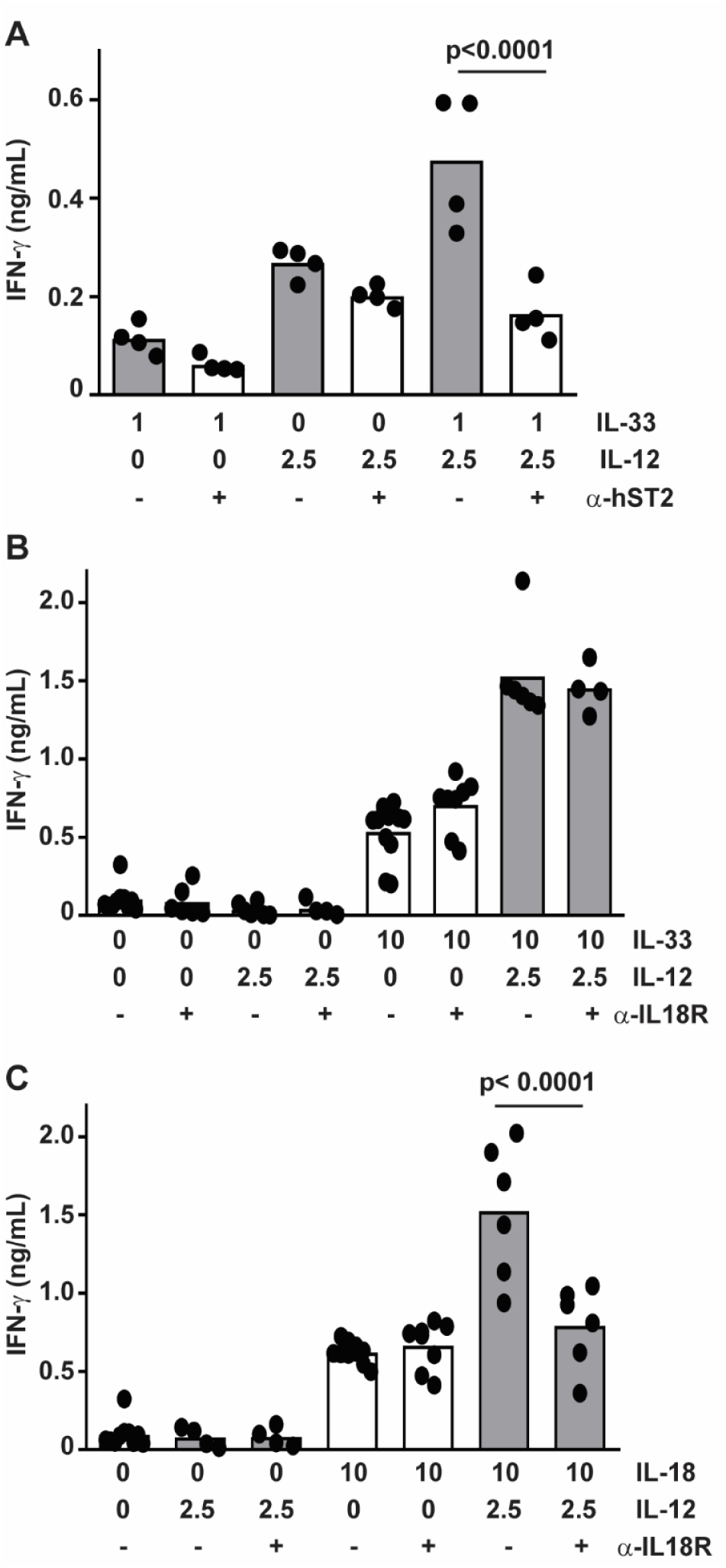
Role of ST2 in IL-33-mediated enhancement of IL-12 responsiveness of NK cells. Enriched primary human NK cells were pre-treated with (**A**) 1 μg anti-ST2 or (**B**,**C**) 0.1 μg anti-IL-18R antibody (open bars) or isotype (closed bars) one hour prior to stimulation with combinations of IL-12 and (**A**,**B**) IL-33 or (**C**) IL-18 (doses listed in ng/mL) for 16 hours prior to ELISA determination of IFN-γ (columns represent mean). Multiple comparisons of ordinary one-way ANOVA were used to compare groups with p<0.1 shown. Each dot represents the mean of three biological replicates of an independent subjects (n=4-7).

### IL-33 enhances IFN-γ and TNF release via activation of p38 MAPK

In macrophages, IL-33 stimulates activation of p38 MAPK^21, 22^. To determine if IL-33 activates p38 MAPK in human NK cells, we measured phosphorylation of p38 MAPK and downstream targets of this kinase by flow cytometry. Primary human NK cells were cultured and stimulated with IL-12, IL-33 or both cytokines. Phosphorylation of p38 MAPK (Thr180, Tyr182) and activating transcription factor 2 (ATF2, Thr71) were measured by flow cytometry. IL-33 stimulation, irrespective of the presence of IL-12, stimulated p38 MAPK phosphorylation within 5 minutes of cytokine exposure (**Fig. 4A**). Levels of phospho-p38 MAPK remained elevated for at least 30 minutes post-stimulation with IL-33. Likewise, IL-33 stimulation induced the phosphorylation of ATF2, albeit with slower kinetics (**Fig. 4B**). In contrast, IL-12 but not IL-33 induced phosphorylation (Tyr693) of STAT4 (**Fig. 5**). The extent of IL-33-induced phospho-p38 MAPK (**Fig. 4A**) and phospho-ATF2 (**Fig. 4B**), as well as IL-12-induced phospho-STAT4 (**Fig. 5**), were not measurably altered over this time scale by co-administration of the other cytokine.

**Figure 4.**
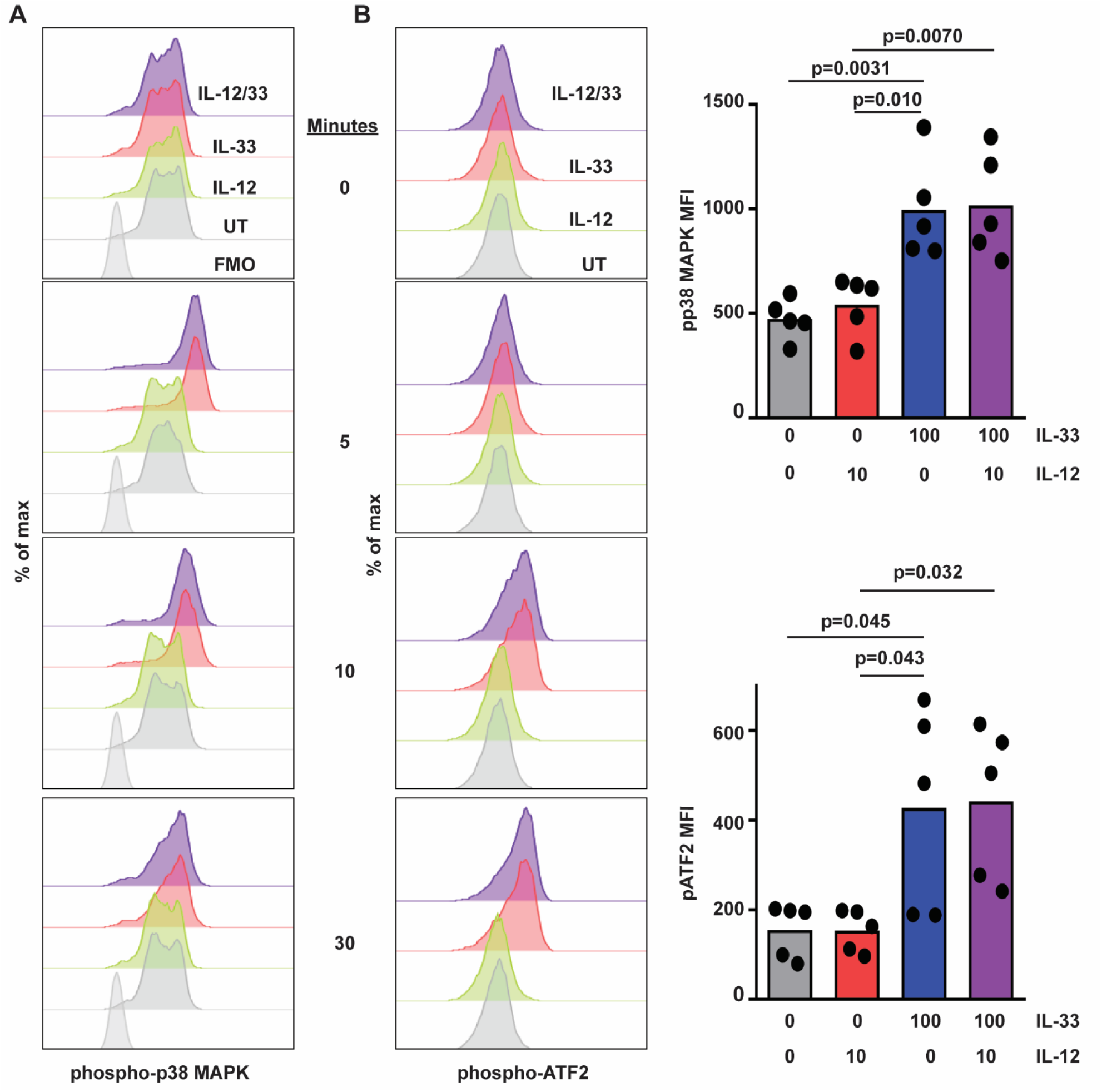
IL-33 activates p38 MAPK in human NK cells. Human NK cells were left untreated (UT) or incubated with IL-12 or IL-33 (doses in ng/mL indicated on right) for 5 to 120 minutes prior to phosphoflow measurement of intracellular **(A)** phospho-p38 MAPK, **(B)** phospho-ATF2 levels. Representative histogram plots are shown (one of five similar experiments with cells from different subjects), along with graphs of median of fluorescence intensity (MFI, each dot represents results obtained from an independent donor). Multiple comparisons of ordinary one-way ANOVA were used to compare columns with p<0.1 shown.

**Figure 5.**
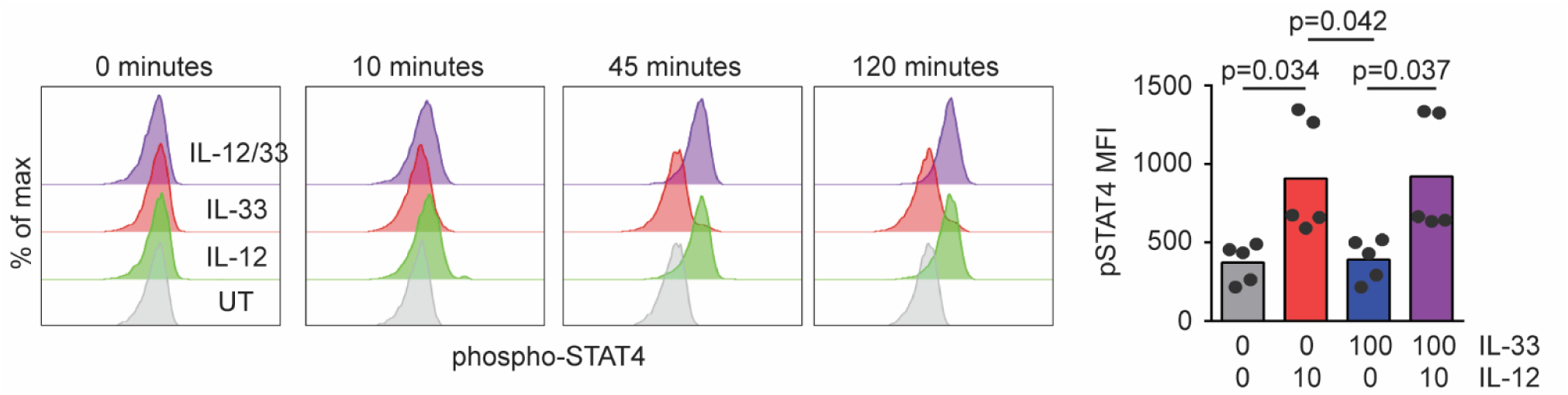
IL-33 does not interfere IL-12 STAT4 phosphorylation. Human NK cells were left untreated (UT) or incubated with IL-12 or IL-33 for 10 to 120 minutes prior to phosphoflow measurement of phosphorylated STAT4 levels. Representative histogram plots are shown (one of five similar experiments using cells from different donors), along with graphs of median fluorescence intensity (MFI, each dot represents results obtained from an independent donor). Multiple comparisons of ordinary one-way ANOVA were used to compare columns with p<0.1 shown.

To determine the role of p38 MAPK in IL-33 enhancement of IL-12-induced cytokines, a specific p38 MAPK inhibitor, R-1503^21^, was employed. NK cells were pre-treated with R-1503 (0 to 500 nM) for 150 minutes prior to stimulation with IL-33 (10 ng/ml) and IL-12 (1 ng/ml) for 16 hours. None of the concentrations of inhibitors used in this study measurably affected cell viability (data not shown). A dose of 125 nM R-1503 restored expression of IFN-γ in response to IL-12 plus IL-33 to levels seen with IL-12 alone (**Fig. 6A**). This concentration of R-1503 did not measurably impact IL-12-induced IFN-γ expression in the absence of IL-33 (**Fig. 6B**), but did diminish the enhancing effect of IL-33 on IL-12-induced TNF expression (**Fig. 6C**).

**Figure 6.**
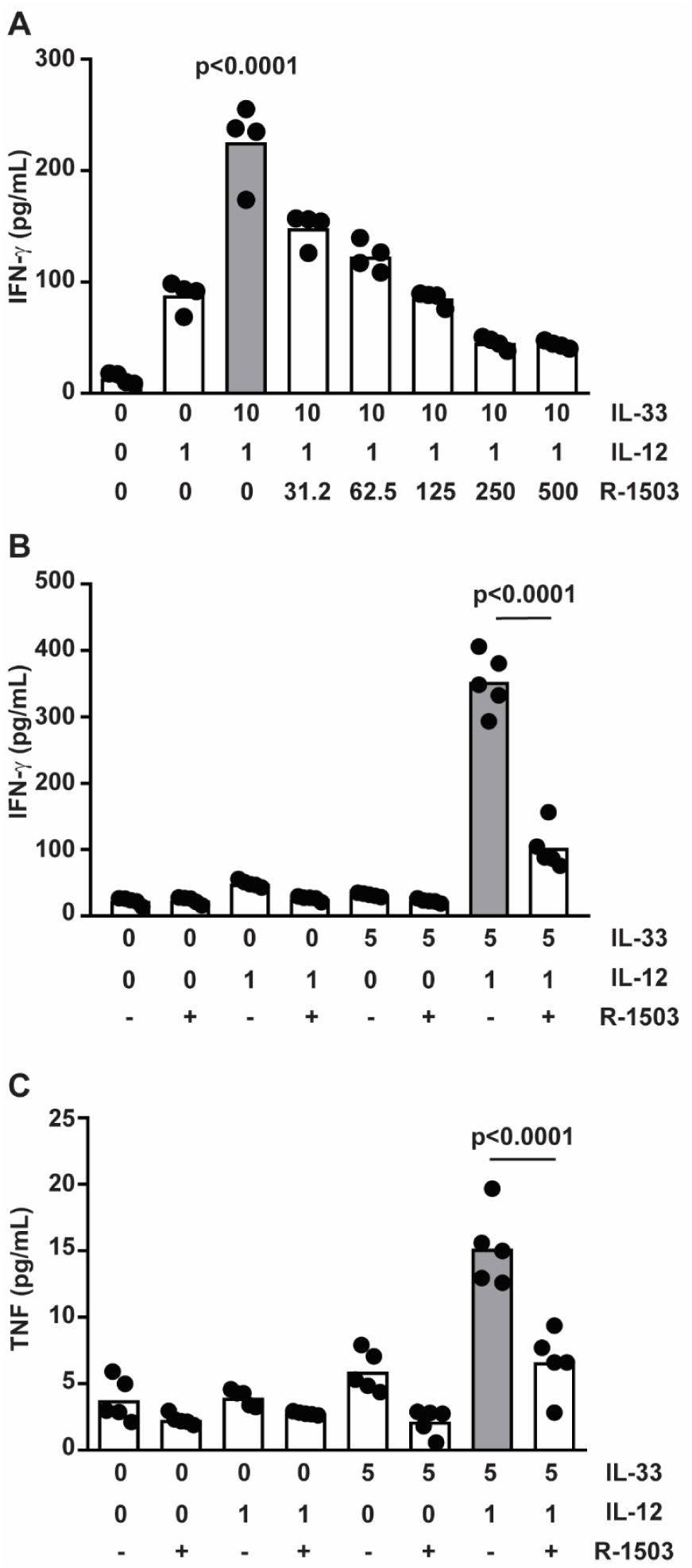
p38 MAPK inhibition abrogates IL-33-enhancement of IL-12-induced activity. **(A)** IFN-γ production (pooled results of 4 independent subjects) by human NK cells following 150-minute pretreatment with vehicle (0.1% DMSO) or various concentrations of R-1503 (0-500 nM) and subsequent stimulation with IL-12 and IL-33 (doses listed in ng/mL). **(B, C)** Effect of pre-treatment with 125 nM R-1503 on levels of IFN-γ and TNF in cell-free supernatant collected 6 hours after cytokine stimulation. Each dot represents the mean of three biological replicats of an independent subject (n=5). Multiple comparisons of ordinary one-way ANOVA were used to compare groups with p<0.1 shown.

IL-18 can enhance IL-12-provoked IFN-γ production by promoting *IFNG* transcript stability, which is mediated by p38 MAPK signaling^23^. To determine whether IL-33 exerts a similar effect on *IFNG* transcript stability, human NK cells were incubated with IL-12 alone or in combination with IL-33 for 6 hours in conjunction with pretreatment with R-1503 or vehicle control. Four hours after cytokine administration, cultures were treated with actinomycin D (5 μg/mL) to halt *de novo* transcription, thereby permitting measurement of decay of existing *IFNG* transcripts. As shown in **Fig. 7**, *IFNG* levels dropped in all experimental groups after addition of actinomycin D. Treatment of NK cells with IL-33 and IL-12 (**Fig. 7**, dark purple) resulted in a significantly greater half-life of *IFNG* transcripts (116±3 minutes for IL-33/IL-12 versus 75±3 minutes for IL-12, p<0.0001) compared to NK cells stimulated with IL-12 alone (**Fig. 7**, solid black lines, dark gray bar). The addition of R-1503 accelerated the decay of *INFG* transcripts (**Fig. 7** left plot, dashed lines), and largely abrogated the enhancing effects of IL-33 (**Fig. 7**, right plot, light purple bar) relative to IL-12 alone (**Fig. 7**, right plot, light grey bar) in terms of half-life of *IFNG* transcripts (62±2 minutes for IL-33/IL-12 versus 52±2 minutes for IL-12, p=0.045). Thus, IL-33 acts to enhance *IFNG* transcript stability via p38 MAPK.

**Figure 7.**
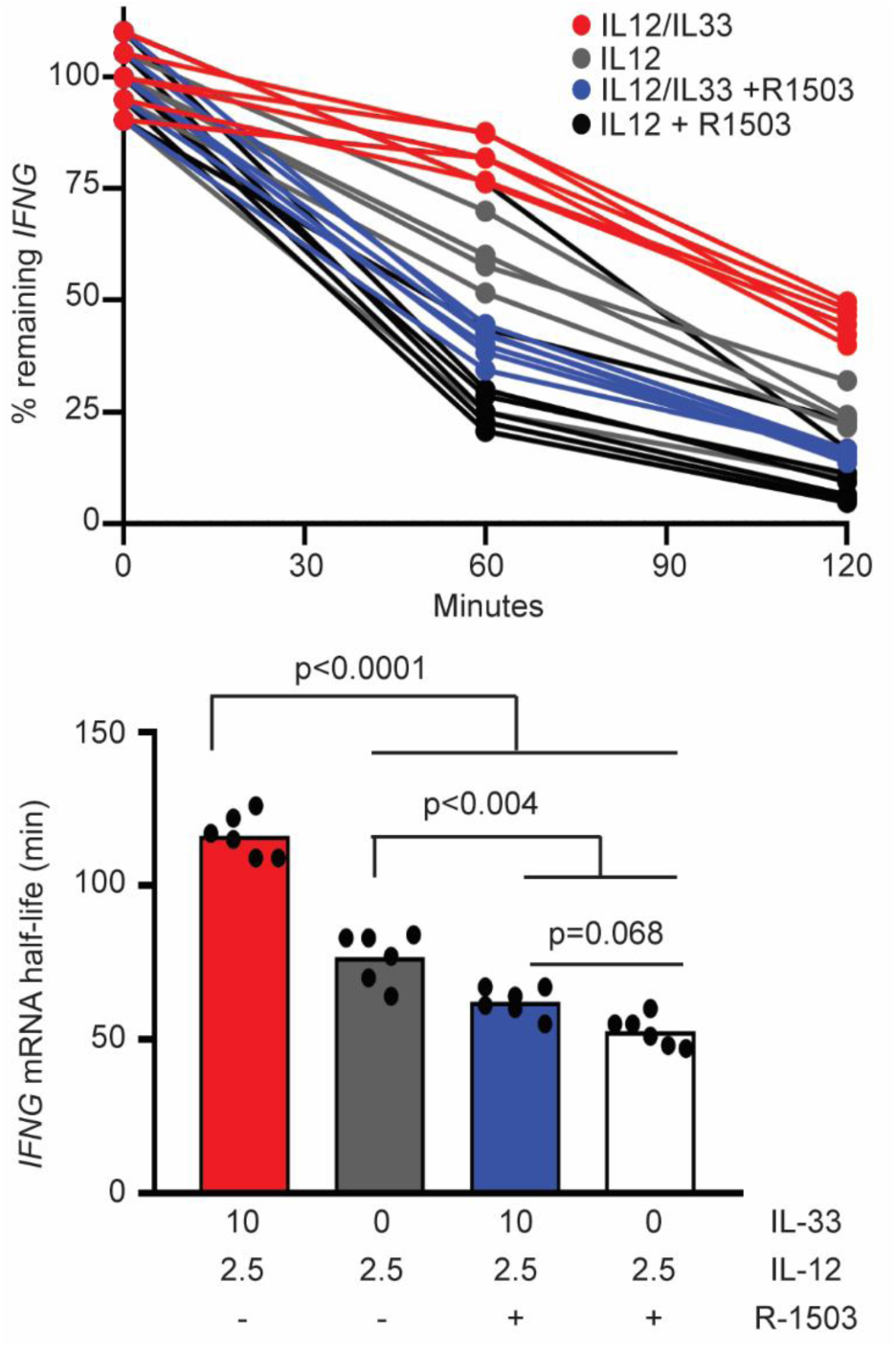
p38 MAPK inhibition disrupts IL-33-mediated IFNG mRNA stability. NK cells were pretreated with R-1503 or DMSO 0.1% and then stimulated with IL-12 (2.5 ng/ml, black lines) or with IL-12/33 (2.5 and 10 ng/ml respectively, purple lines) for 4 hours. After 4 hours, Actinomycin D (ActD 5 µg/ml, solid lines) or Actinomycin D with R-1503 (125 nM, dashed lines) were added. *IFNG* expression was measured by qRT-PCR (left) and *IFNG* mRNA half-life was calculated (right) via linear regression. Each line or dot represents one of two biological replicates for one of three different subjects. Multiple comparisons of ordinary one-way ANOVA were used to compare cell treatments.

### MAPKMAPK (MK2) contributes to IL-33 enhancement of IL-12-induced IFN-γ

Previous studies showed that MAPKMAPK (MK2) is down-stream of p38 MAPK, and that MK2 mediates inflammatory responses^24, 25^. Addition of a specific MK2 inhibitor (MK2 IV) to NK-cell cultures diminished the enhancing effect of IL-33 on IL-12-stimulated IFN-γ production (**Fig. 8**). In contrast to R-1503, no dose of MK2 IV that we tested completely abrogated IL-33 enhancement of IFN-γ expression. Thus, MK2 contributes to the enhancing effect of IL-33 but does not fully account for the effects mediated by p38 MAPK.

**Figure 8.**
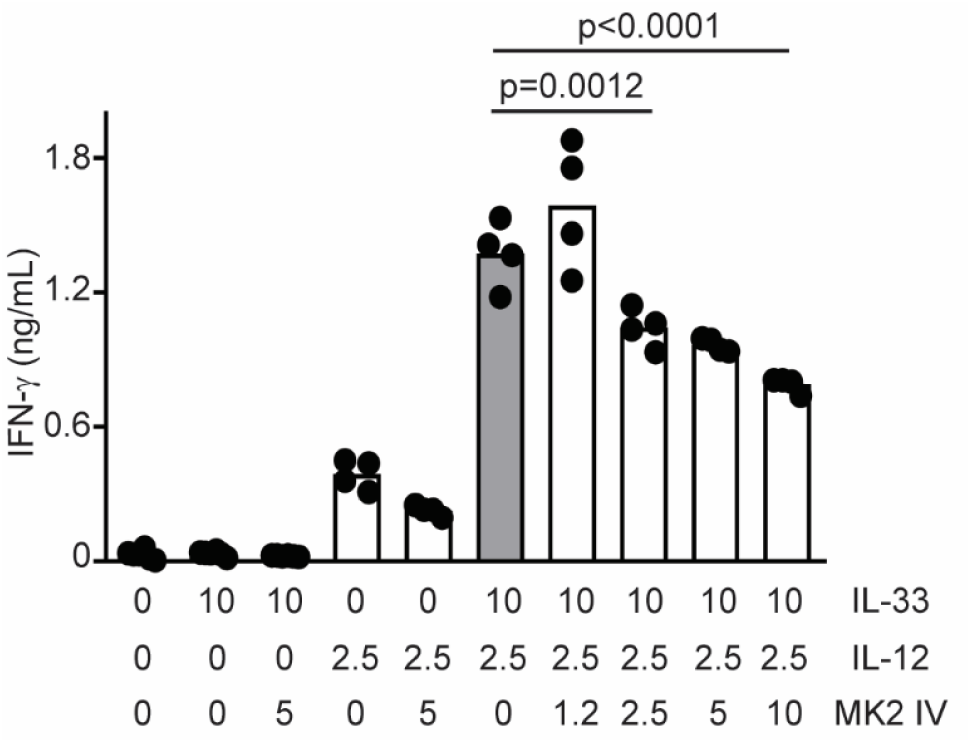
MK2 inhibition interferes IL-33 enhancement of IL-12 induced activity. IFN-γ production by human NK cells following 150-minute pretreatment with vehicle (0.1% DMSO) or various concentrations of MK2 IV (0-10 µM) and subsequent stimulation with IL-12 (2.5 ng/mL) and IL-33 (10 ng/mL) for 16 hours, IFN-γ levels were measured in cell free supernatant. Each dot represents the mean of three biological replicates of an independent subject (n=4). Multiple comparisons of ordinary one-way ANOVA were used to compare columns with p<0.1 shown.

### IL-33 and IL-12 regulate TNF release via ADAM17

TNF is shed from the cell membrane via activity of ADAM-17^26^. In addition to IL-33 induced increases in *TNF* mRNA expression (**Fig. 1B**), we hypothesize that IL-33 further enhances TNF secretion by NK cells (**Fig. 2B**) via elevated ADAM-17 cleavage of cell-surface TNF. Addition of the ADAM17 inhibitor (TAPI-1)^27^ to NK-cell cultures resulted in reduction of secreted TNF from 97.1±10.4 to 45.8±5.2 pg/ml (ELISA) in response to IL-33 and IL-12 stimulation (**Fig. 9A**). In contrast, R-1503 more completely ablated the enhancing effect of IL-33 on IL-12-induced TNF secretion. Thus, enhanced TNF secretion in response to IL-12 and IL-33 is at least partially dependent on ADAM-17, which may function downstream of p38 MAPK or be dependent on p38 MAPK induced changes in *TNF* mRNA expression. While secreted TNF secretion increased in response to the combination of IL-33 and IL-12, cell-associated TNF was reduced 1.9-fold to levels near those induced by IL-12 alone (**Fig. 9B**). These flow cytometry measurements were made in the absence of brefeldin-A or other secretion inhibitors, thus reflecting TNF naturally retained by the cells. Importantly, reduced TNF secretion by TAPI-I-treated, IL-12 and IL-33 stimulated cells (**Fig. 9A**) was associated with a 2.0-fold increase in cell-associated TNF to levels observed with IL-12 alone (**Fig. 9B**). Thus, IL-33-stimulated ADAM17 activity contributes to TNF secretion at the expense of cell-associated TNF, likely via cleavage of TNF from the cell surface.

**Figure 9.**
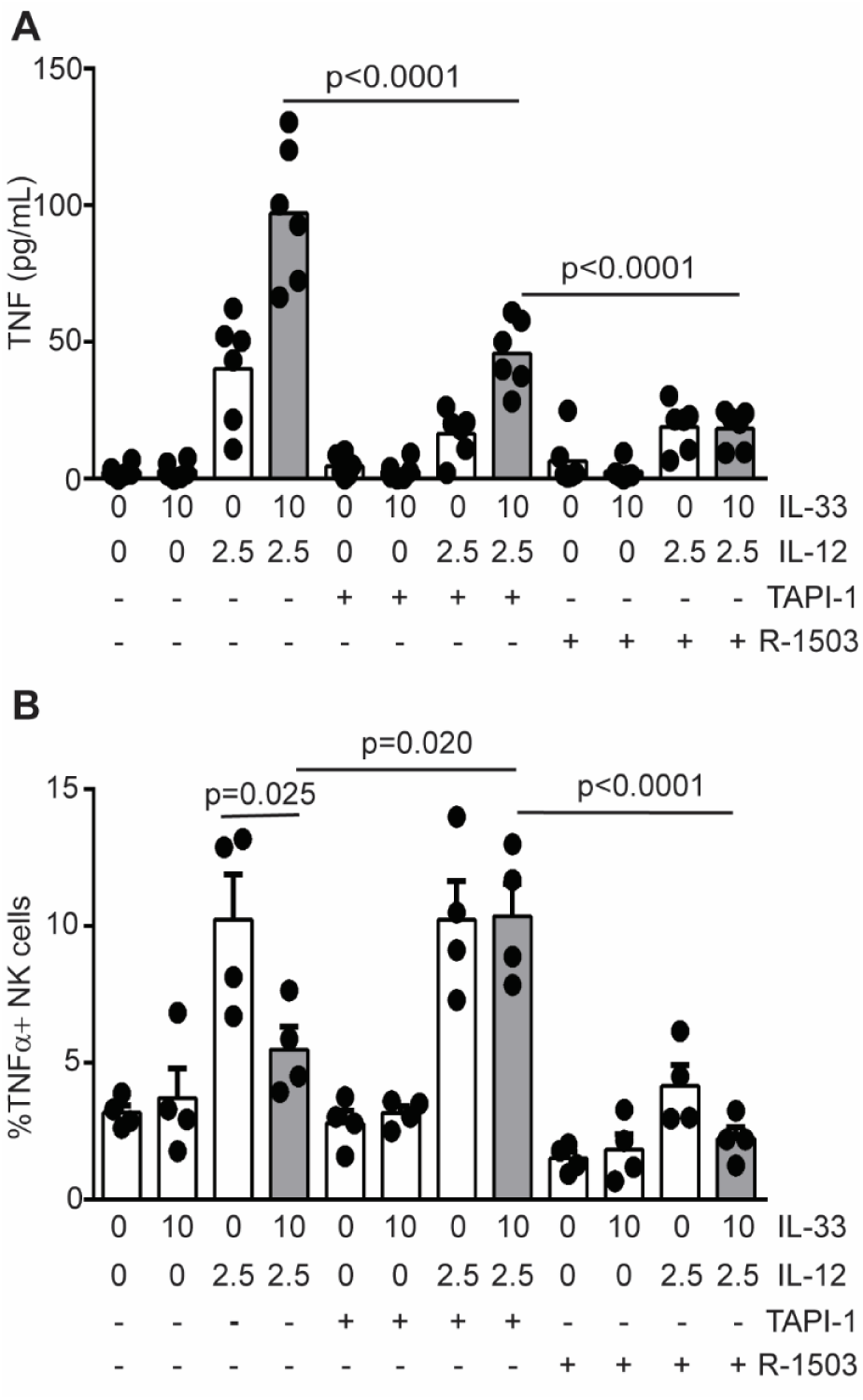
IL-33 and IL-12 regulate TNF release via ADAM17. Enriched human NK cells were pretreated with 0.1% DMSO, R-1503 (0.25 µM) or TAPI-1 (8 µM) and then stimulated in the presence of cytokines (doses listed in ng/mL) for 16 hours. (**A**) Secreted TNF expression was measured by ELISA where each dot represents the mean of three biological replicates of an independent subject (n=6). **(B)** Cell-associated TNF was measured by flow cytometry where each dot represents the mean of two biological replicates of an independent subject (n=4). Multiple comparisons of ordinary one-way ANOVA were used to compare columns with p<0.1 shown.

## Discussion

We augment previous observations in mouse^17, 18^ and human^16^ NK cells by showing that near-physiological (e.g. pM) concentrations^28, 29^ of the type 2 cytokine IL-33 trigger hypersensitivity of primary human NK cells to the type I cytokine IL-12. In addition, superphysiological concentrations of IL-33 and IL-12 synergistically induced the release of TNF and GM-CSF, suggesting that high concentrations of these cytokines in a tissue microenvironment may have even stronger effects on NK-cell function. IL-33-mediated potentiation of IFN-γ and TNF secretion depended on p38 MAPK, and to a lesser extent, the downstream kinase MK2. Moreover, enhanced TNF secretion was partially attributable to ADAM17. These results highlight the possibility for exaggerated cytokine secretion by human NK cells in physiological setting that are characterized by dual exposure to type 1 (IL-12) and type 2 (IL-33) cytokines.

The type 1 cytokine, IL-18, enhances expression of *IFNG* by IL-12-stimulated human NK cells^30^ in part via p38 MAPK stabilization of *IFNG* transcripts^23^. Our data suggest that a type 2 cytokine, IL-33, similarly enhances IL-12-induced IFN-γ expression by human NK cells. Moreover, IL-33 amplification of IFN-γ is dependent on p38 MAPK activity and associated with increased stability of *IFNG* mRNA. The p38 MAPK regulates inflammatory responses via phosphorylation of downstream mediators, including MK2^31, 32^. Yet, MK2 inhibitors had only a partial inhibitory effect in contrast to the abrogation of IL-12/IL-33 synergistic interactions following p38 MAPK inhibition. This finding implies that other p38 MAPK-stimulated mediators, including ATF2, may also be crucial for the enhancing effect of IL-33. In fact, IL-33 rapidly induced the phosphorylation of p38 MAPK and ATF2 in human NK cells. Of note, p38 MAPK-induced activation of ATF2 initiates activator protein 1 (AP-1) complex-mediated transcription of inflammatory cytokines, including IFN-γ ^33-35^. Coupled with the lack of apparent effects of IL-33 on IL-12-induced phosphorylation of STAT4, our results highlight the possibility that IL-33 enhances IFN-γ via enhanced AP-1-dependent promotion of transcription.

In addition to p38 MAPK-driven intracellular signaling leading to transcription factor activation, p38 MAPK can also directly phosphorylate membrane proteins like ADAM17^36^. The activation of ADAM17 by p38 MAPK can promote shedding of TNF^21, 37^. Our data are consistent with these possibilities, revealing p38 MAPK-dependent enhancement of TNF expression coupled to ADAM17-dependent release of cell-associated TNF. Collectively, our data suggest that p38 MAPK govern several molecular pathways which regulates inflammatory cytokine release in human NK cells.

The ability of type 2 cytokines to mediate IFN-γ release is not limited to IL-33. Past studies demonstrated that IL-4 enhance IFN-γ production in murine IL-12 and IL-15 stimulated NK cells^15, 38^. We also observed IL-4 or IL-13 enhancement of IL-12-induced IFN-γ production by human NK cells (D.O. and S.N.W. unpublished observations). Whereas IL-4 and IL-13 predominately signal via activation of STAT6, both cytokines appear capable of triggering the p38 MAPK pathway^39^. In fact, synergy between IL-4 and IL-12 in mouse NK cells is partially dependent on p38 MAPK^15^. Whether distinct type 2 cytokines enhance IL-12-induced IFN-γ production via similar or distinct mechanisms remains to be determined.

In pathologies such as asthma and COPD, IL-33 play a central role in disease pathology, those pathologies were shown to be exacerbated by respiratory viral infection which mediates the release of type 1 cytokines such as IL-12^40-43^. Thus, we speculate that NK cells derived from the blood or tissues of patients exhibiting elevated levels of IL-33 would be more sensitive to *ex vivo* IL-12 stimulation. As an example, cigarette smoke induces epithelial damage resulting in increased expression of IL-33^17, 42^, which provokes hypersensitive IFN-γ production by mouse NK cells in response to IL-12^44^. Our initial data suggest that NK cells from smokers exhibit greater sensitivity to IL-12 in terms of IFN-γ expression than NK cells derived from healthy, non-smokers (D.O., M.B., and S.N.W unpublished observations). We speculate that elevated IL-33 levels in smokers or asthmatic patients provokes hypersensitivity to type 1 inflammatory cues triggered by virus infection. Thus, smokers exposed to virus infections likely produce high, potentially pathogenic expression of IFN-γ. Our findings provide new insights about potential functional hypersensitivity of NK cells in type 2 cytokine rich inflammatory milieus such as asthma and COPD as well as in smokers.

## Materials and Methods

### Human cells

De-identified blood samples were obtained from healthy donors, as defined by Hoxworth Blood Center guidelines (https://hoxworth.org/donors/eligibility.html), with the approval of Cincinnati Children’s Hospital Medical Center Institutional Review Board. Peripheral blood mononuclear (PBMCs) cells were isolated on a ficoll-hystopaque (GE, Marlborough, MA) gradient, and NK cells were enriched by negative selection using the NK-cell isolation kit immune magnetic beads per manufacturer’s protocol (Miltentyi Biotec, San Diego, CA). Enriched cells were >90% CD56^+^ CD3^neg^ NK cells as determined by flow cytometry.

### In vitro culture of NK cells

Enriched NK cells were cultured in SCGM media (Cellgenix®, Freiburg, Germany) supplemented with IL-2 (400 U/ml, Peprotech, Rocky Hill, NJ), 10% human serum (Sigma, St. Louis, MO), 10% heat-inactivated fetal bovine serum, 100 U/ml penicillin, 100 U/ml streptomycin, 1% non-essential amino acids, 1% sodium pyruvate, 2mM L-Glutamine and 10 mM HEPES. In experiments, NK cells were either used directly after blood isolation or after 7 to 30 days of *ex vivo* culture. Each scenario produced similar results.

### Antibodies

The following antibodies were used in the described studies: Brilliant violet (BV) 421-CD56 (5.1H11), BV711-CD16 (3G8), AlexaFluor (AF) 647-NKp46 (9E2), PE-NKp44 (P44-8) were purchased from Biolegend (San Diego, CA); Brilliant Ultra violet (BUV) 395-CD3 (SK7), V500-CD3 (SP34-2), BUV797-CD69 (FN50), and BUV-CD62L (SK11) were purchased from Becton and Dickinson (San Diego, CA); eFluor710-NKG2D (1D11) from Invitrogen (Carlsbad, CA); and mouse anti-human ST2 (AF523 or unconjugated) and anti-human IL-18R (70625) from R&D (Minneapolis, MN). Antibodies were used at doses titrated in our lab.

### Cytokine stimulation

Enriched NK cells (unless mentioned otherwise, 5×10^4^ per well in triplicate per condition) were cultured overnight in the presence of 60 U/ml of IL-2. Various concentrations of IL-12, IL-18 or IL-33 or (Peprotech®, Rocky Hill, NJ) were added to cultures for 6 to 16 hours. Cell-free supernatant was collected and analyzed by ELISA for levels of human IFN-γ, TNF and GM-CSF (Invitrogen, Waltham, MA). For inhibition of mitogen-activated protein kinase (MAPK) activity, 125 nM (measured IC50) of the selective p38 MAPK inhibitor Pamapimod (R-1503, Selleckchem, Houston, TX) was added 150 minutes prior to cytokine stimulation. The inhibition of MK2 (p38/mitogen-activated protein kinase-activated protein kinase 2) was performed with 5 μM MK2 IV (CAYMAN chemical, Ann Arbor, MI). The inhibition of ADAM-17 (a disintegrin and metalloprotease-17) was performed by adding 8 μM TAPI-1 (Merck Millipore, Burlington, MA). TAPI-1 and MK2 IV were added 60 minutes before cytokine stimulation. At the tested concentrations, none of the tested inhibitor affected cell viability (data not shown).

### mRNA stability assay

Primary human NK cells were cultured at 1×10^5^ cell per well. Cells were treated with media (0.1% DMSO), IL-12 (2.5 ng/ml) with or without IL-33 (10 ng/ml). Actinomycin D (5 μg/mL, Sigma) was added to cell cultures 4 hours after cytokine treatment, and RNA was isolated before addition of actinomycin D and 2 hours after addition of actinomycin D. RNA was extracted from NK cells with RNeasy kit (Qiagen, Hilden, Germany) according to manufacturer’s instructions. The detection of *IFNG, TNF, IL5* and *IL13* mRNA expression was performed by using TaqMan® probes (Applied Biosystems, Foster City, CA) according to manufacturer’s instructions.

### Statistical analysis

We performed statistical analyses using GraphPad Prism 8.01. We used two-way ANOVA to identify the contribution to multiple variables to an experimental measurement. We used one-way ANOVA to perform multiple comparisons between experimental conditions. The specific statistical analysis test used is indicated in each figure legend.

## Author contribution

DEO: conceptualization, investigation, data curation, writing-original draft. AA, PCA, and DK: investigation, data curation, writing-editing. LCK: data curation, writing-editing. MB: data curation, writing-editing. SNW: conceptualization, data curation, writing-original draft, study supervision.

## Acknowledgements

This project was supported in part by NIH P30 DK078392 (Gene Analysis Core) of the Digestive Diseases Research Core Center in Cincinnati and the NIH P30 AR070549 Center for Rheumatic Disease Research. This work was supported by NIH grants DK107502 (L.C.K.), AR073228 (S.N.W. and L.C.K.), HL1195338 (M.T.B.), HL141236 (M.T.B.), AI148080 (S.N.W.), and DA038017 (S.N.W.); the Veteran’s Administration (I01BX002347 to MTB); research initiation funds (S.N.W. and L.C.K.) and an Arnold W. Strauss Fellow Award (D.E.O.) from the Cincinnati Children’s Research Foundation; and a postdoctoral fellowship form the American Heart Association (D.E.O.). A.A and P.C.A were supported by MSTP NIH T32 GM063483, while A.A. is also supported by T32 AI118697.

## Notes

#### Summary of Updates

Changes made to text and several figures in response to peer review. New data in Figure 1C as well as Figure 3B-C, and revised analysis/presentation of Figure 7.

